# Artificial Brain Extracellular Fluid in Radioligand Binding Assays: Effects on Affinities of Ligands at Serotonin 5-HT7 Receptors

**DOI:** 10.1101/2025.09.04.674239

**Authors:** Anh T. Huynh, Hamad M. Alali, Jason V. Wallach, Clinton E. Canal

## Abstract

Radioligands are well-established tools for measuring ligand binding affinities at receptors. Determining affinities of test ligands at many G protein-coupled receptors (GPCRs), including serotonin (5-HT) GPCRs, often involves incubating a radioligand, test ligands, and receptors expressed in cell membranes in Tris buffers, and commonly in a standard binding buffer (SBB) containing Tris HCl, MgCl_2_, and EDTA until ligand– receptor equilibrium binding is established. However, the composition of extracellular fluid (ECF), where ligands first encounter GPCRs in vivo, differs from that of SBB, which we hypothesized impacts ligand affinity. We conducted radioligand binding assays to compare the affinities of the agonist 5-carboxamidotryptamine (5-CT) and two antagonists/inverse agonists, lurasidone and SB-269970, at [^3^H]5-CT-labeled 5-HT7 GPCRs stably expressed in HEK293 cells using SBB or artificial brain ECF (abECF) as the medium at room or physiological temperatures (RT or 37°C). The rank order of ligand potencies, as well as 5-CT’s affinity, was unaffected by the different experimental environments. [^3^H]5-CT 5-HT7R B_max_ values increased in abECF and modestly at 37°C, without affecting *K*_d_, suggesting an increase in active state conformations. In contrast to 5-CT, antagonist/inverse agonist affinities depended on both media and temperature. The affinities of lurasidone and SB-269970 at 5-HT7 receptors were substantially higher at 37°C than at RT. Also, incubation of lurasidone and SB-269970 in abECF resulted in significantly higher affinities compared to incubation in SBB (e.g., ∼10-fold higher for lurasidone), indicating that temperature and the buffer and ionic composition of abECF influence 5-HT7 antagonist/inverse agonist ligand binding. As a high concentration of NaCl in abECF is a remarkable difference from the composition of SBB, we probed the impact of removing NaCl from abECF; removal of NaCl had a minor affinity-enhancing effect on the antagonists, inferring that other ions, glucose, or sodium phosphates in abECF underlie significant changes in ligand–receptor binding interactions. Overall, the observations indicate that measuring 5-HT7 antagonist affinities at [^3^H]5-CT-labeled 5-HT7Rs with abECF at physiological temperature—modeling the in vivo brain environment where ligands and GPCRs interact—yields distinct affinity values that may be more physiologically accurate than values obtained from SBB. Moreover, several historical reports demonstrate that temperature, ions, and buffers have no consistent effect on the affinities of distinct ligands at various other GPCRs, and there is no consensus binding buffer used in the literature for any GPCR, which may contribute to the variability in ligand–GPCR affinities reported. These findings show that buffer and temperature impacted 5-HT7R ligand binding affinities and highlight the importance of considering such conditions when performing experiments.

## Introduction

Ligand–G protein-coupled receptor (GPCR) binding is fundamental to initiate numerous biological functions. As endogenous ligands bind to their cognate GPCRs, typically expressed in cell membranes, they stimulate intracellular G protein signaling cascades that precisely modulate cellular responses, e.g., smooth or cardiac muscle cell contraction, glandular epithelial cell secretion, or neuronal action potential firing. GPCRs are also one of the most prominent drug target families, representing 36% of all FDA-approved medicines (Lorente et al., 2025). Medications that bind to GPCRs can potentiate or inhibit their natural behaviors in various ways, leading to therapeutic outcomes by supporting cellular, tissue, or organ homeostasis. Determining ligand affinities and functional potencies at GPCRs in biological systems is vital for drug discovery, as they inform decisions regarding physiologically relevant target engagement, compound efficacy, and structure–activity relationships. Also, many GPCR-targeting drugs—especially drugs to treat central nervous system (CNS) disorders—interact with multiple GPCRs, but their overall contributions to therapeutic efficacy and side effects are poorly understood, creating a challenge from the start to fine-tune polypharmacology for improved outcomes (Du et al., 2016; Kabir and Muth, 2022; Kenakin, 2019; Roche et al., 2020; Roth et al., 2004).

Most ligands reversibly bind to GPCRs. A common technique for determining the binding affinity of a test agent with reversible binding at GPCRs is the radioligand competition binding assay (Auld, 2012; Kenakin, 2019; St John-Campbell and Bhalay, 2025). By measuring the competition of a test ligand and a high-affinity radioligand for receptor binding sites, one can determine the equilibrium inhibitory constant, *K*_i_, of the test ligand, which is the mathematical affinity of the ligand at the receptor that incorporates the concentration of the ligand required to inhibit 50% of binding of the radioligand, the radioligand’s affinity (*K*_d_) at the receptor, and the radioligand’s concentration in the assay (Yung-Chi and Prusoff, 1973). *K*_i_ values are often expressed on a logarithmic scale as p*K*_i_, as the concentrations of test ligands are reported graphically on a logarithmic scale; concentration parameters usually have a log-normal distribution, making standard deviations symmetric for p*K*_i_ values, but not for *K*_i_ values (Christopoulos, 1998). Radioligand saturation binding assays, on the other hand, are used to determine the radioligand’s affinity, *K*_d_, or equilibrium dissociation constant, the concentration of the ligand at which half of the receptors are occupied at equilibrium, as well as the total number of receptor binding sites, B_max_, in the protein test samples.

Although radioligand binding assays have been a mainstay in pharmacology for decades, the experimental conditions for performing them are often inconsistent across receptors, ligands, and independent laboratories. However, many incubation buffers contain Tris (hydroxymethyl)aminomethane (Tris) buffer because it has a physiologically relevant, wide buffering range and is versatile for molecular biology applications. A widely used buffer, including by our lab and collaborators, is the standard binding buffer (SBB) that contains 50 mM Tris-HCl, 10 mM MgCl_2_, and 0.1 mM EDTA. Yet, Tris is a non-natural buffer, and SBB lacks various ionic compounds in the in vivo extracellular milieu, where typical ligands bind to their binding sites and encounter the target GPCRs expressed in the cell membrane. Suppose ligands have different affinities when tested in SBB than in physiological media. In that case, affinities derived from incubating ligands in SBB have the potential to misguide structure–activity relationship studies, the results of which inform decisions in the drug discovery process, and without careful target engagement studies in vivo, could misguide considerations about the contributions of distinct GPCRs to a drug’s pharmacodynamic effects (Auld, 2012; St John-Campbell and Bhalay, 2025; Sum et al., 2019). This would become especially relevant when comparing binding affinities across GPCRs or other target proteins to determine a ligand’s selectivity profile.

This project focused on establishing an environment for radioligand binding assays that mimics the in vivo extracellular brain milieu, where CNS drugs bind to receptors expressed in brain cells. We compared affinities of ligands at the 5-hydroxytryptamine (serotonin, 5-HT) 7 receptor (5-HT7R) incubated in SBB versus an artificial brain extracellular fluid (abECF), the contents of which were determined from chemical analysis of small molecules in the extracellular fluid collected from the hippocampus (McNay and Sherwin, 2004). We conducted these radioligand binding assays at both room temperature (RT) and physiological temperature (37°C), as temperature can influence the accessibility of GPCR binding sites and impact ligand affinity (Auld, 2012; Hall et al., 1990; Pasternak and Pan, 2013).

The 5-HT7R—expressed as two functionally similar isoforms, 5-HT7Ra and 5-HT7Rb (Jasper et al., 1997)— was the last 5-HTR subtype cloned and is present in the CNS and periphery. In the CNS, 5-HT7Rs are expressed in the hypothalamus, thalamus, and hippocampus (Horisawa et al., 2013), and can impact circadian rhythms, sleep, and memory (David and Jim, 2004). In the periphery, 5-HT7Rs regulate blood pressure, with activation promoting vasodilation (Jackson et al., 2023). 5-HT7Rs are also putative targets for the treatment of chronic schizophrenia and bipolar disorder symptoms, and lurasidone (Latuda®), an antipsychotic drug, possesses polypharmacology that includes potent 5-HT7R antagonism (Nikiforuk, 2015; Okubo et al., 2021; Roth et al., 1994; Teitler et al., 2010; Wei et al., 2020). We measured the affinity of three 5-HT7R ligands: 5-carboxyamidotryptamine (5-CT), a potent full agonist (Armstrong et al., 2020; Bard et al., 1993; Brüss et al., 2005); SB-299670, a potent antagonist/inverse agonist (Mahé et al., 2004; Romero et al., 2006); and lurasidone, a potent antagonist/inverse agonist (Ishibashi et al., 2010). Using [^3^H]5-CT as the radioligand, the reported affinities of 5-CT and SB-269970 at the 5-HT7R are p*K*i 9.1 (*K*_i_ ∼0.8 nM) (Thomas et al., 1998) and 8.9 (*K*_i_ ∼1.3 nM) (Hagan et al., 2000), respectively; yet, results were determined using cell membranes expressing human 5-HT7aRs and ligands incubated in 50 mM Tris HCl, 4 mM CaCl_2_, 0.1 mM pargyline (a monoamine oxidase inhibitor), and 1 mM ascorbic acid at pH 7.4 and 37°C for 60 min. Lurasidone also had potent binding affinity at [^3^H]5-CT-labeled 5-HT7Rs with a reported p*K*_i_ of 9.3 (*K*_i_ ∼0.5 nM) (Ishibashi et al., 2010); in this study, the incubation buffer used was 50 mM Tris, 0.5 mM Na_2_–EDTA, 10 mM MgSO_4_, 2 mM CaCl_2_, 0.01 mM pargyline, and 0.1% ascorbic acid (To et al., 1995) at pH 7.4 at RT for 120 min. In another study where [^3^H]SB-269970 was used as the radioligand, the p*K*_i_ of lurasidone was 8.68 (*K*_i_ ∼ 2.1 nM) (Horisawa et al., 2013); the incubation buffer used was 50 mM Tris–HCl (pH 7.4), 4 mM CaCl2, 0.1 mM pargyline, and 1 mM ascorbic acid at 37°C for 60 min.

We hypothesized that 5-HT7R ligand affinities would differ when using abECF compared to SBB and that, since abECF mimics the native extracellular fluid composition of the brain, such findings would encourage greater focus on buffer selection and conditions for competitive radioligand binding experiments.

## Materials and Methods

### 1. Buffer Preparation

#### 1.1. Standard Binding Buffer (SBB)

SBB consisted of 50 mM Tris-HCl, 10 mM MgCl_2_, and 0.1 mM EDTA, at pH 7.4 (**Table 1**). Tris-HCl and Tris base were purchased from Fisher Scientific. MgCl_2_ and EDTA were purchased from Sigma Aldrich and Invitrogen. After preparation at room temperature (RT), the SBB was sterilized through 0.22 μm filtration, aliquoted, and stored at 4°C before use. On test days, after bringing SBB to RT, the pH was confirmed to be ∼7.4 before use. After bringing separate aliquots of SBB to 37°C, the pH was adjusted to ∼7.4 with HCl and NaOH. Concentrations of test ligands were then prepared in the SBB solutions.

**Table 1.** Compositions of SBB and abECF. The pH of SBB and abECF was set at 7.4 before each assay.

| SBB | abECF |
| --- | --- |
| Tris HCl: 50 mM | NaCl: 128 mM |
| MgCl<sub>2</sub>: 10 mM | KCl: 3.0 mM |
| EDTA: 0.1 mM | CaCl<sub>2</sub>: 1.3 mM |
|  | MgCl<sub>2</sub>: 1.0 mM |
|  | Na<sub>2</sub>HPO<sub>4</sub>: 21.0 mM |
|  | NaH<sub>2</sub>PO<sub>4</sub>: 1.3 mM |
|  | Glucose: 1.06 mM |

#### 1.2. Artificial Brain Extracellular Fluid (abECF)

abECF consisted of 128 mM NaCl, 3.0 mM KCl, 1.3 mM CaCl_2_, 1.0 mM MgCl_2_, 21.0 mM Na_2_HPO_4_, 1.3 mM NaH_2_PO_4_, and 1.26 mM D-(+)-Glucose, at pH 7.4. NaCl, KCl, Na_2_HPO_4_, and NaH_2_PO_4_ were purchased from Fisher Scientific, and CaCl_2_ and D-(+)-Glucose were purchased from Sigma Aldrich. The solution, prepared at RT, was sterilized via 0.22 μm filtration, and stored at 4°C. After bringing the abECF to RT or 37°C, the pH was adjusted to ∼7.4 with HCl and NaOH before use. Concentrations of test ligands were then prepared in the abECF solutions.

### 2. Cell Culture

Human embryonic kidney cells (HEK293, ATCC, CRL1573) stably expressing the human serotonin 5-HT7aR (Canal et al., 2015) were cultured in Dulbecco’s Modified Eagle Medium with high glucose, 10% dialyzed fetal bovine serum from Gibco, with 500 μg/mL of G418 sulfate from Corning. The cells were seeded in 10 cm Celltreat Petri dishes and grown in an incubator at 37°C, 5% CO_2_, and 95% humidity. The culture medium was replenished every 2 days to maintain a fresh supply of nutrients and to remove metabolic waste. Cells were monitored daily. Cells were passaged when they reached 80% confluency, which involved detaching the cells from the dish surface by the force of pipetting their growth media, counting them, and seeding the cells into new dishes to ensure consistent growth rates.

### 3. Membrane Collection

Cell membranes were collected from HEK293 cultures upon reaching 90% confluency. The cells were gently detached from the plate by the force of pipetting. Once cells were in suspension, they were transferred into 50 mL centrifuge tubes. The cell suspensions were centrifuged at 13,000 x *g* for 10 min at 3°C. Following centrifugation, the supernatant was carefully aspirated. The isolated cell pellets were then resuspended in ice-cold 50 mM Tris-HCl buffer at pH 7.4 and homogenized using a sonicator. This homogenization and centrifugation process was repeated 3 times. Following the final centrifugation, the supernatant was decanted, and the purified cell pellets were resuspended in either SBB or abECF. Cell membranes were aliquoted and stored at -80°C prior to testing.

### 4. Determination of [^3^H]5-CT’s K_d_ and B_max_ under different experimental conditions

Radioligand saturation binding assays were conducted to determine the dissociation constant (*K*_d_) of [^3^H]5-CT at 5-HT7aRs and to evaluate the total receptor binding sites (B_max_) labeled by [^3^H]5-CT in the prepared membranes. Cell membranes (50 µL in buffer) were incubated with concentrations of [^3^H]5-CT ranging from 0.03 nM to 4 nM in SBB or abECF at RT or 37°C in 96-well assay plates (Celltreat, round well, V-bottom). A concentration of 100 μM lurasidone was used to define nonspecific binding. Plates were sealed with transparent tape, wrapped in aluminum foil, and incubated at either RT or 37°C for 90 min. Following incubation, plate contents were vacuumed through GF/B fiberglass filter mats. The filters were then washed with 250 mL of ice-cold, 50 mM Tris-HCl assay buffer (50 mM Tris-HCl, pH 7.4 at 4°C). The wash was repeated twice, and then the filters were dried on a hot plate. The dry filters were then soaked in plastic bags with Betaplate scintillation fluid (PerkinElmer). Bags were sealed and placed in cassettes. Tritium-elicited scintillations were detected and recorded by a Microbeta2 microplate counter (PerkinElmer, tritium efficiency 48%), and disintegrations per minute (DPM), corresponding to each well on the plate, were recorded for two minutes per sample. Two biological replicates, each with three technical replicates per concentration, were tested for each environmental condition. Protein concentrations from membrane preparations were determined using the bicinchoninic acid method (Pierce BCA Protein Assay Kit).

### 5. Determination of ligand K_i_ values at [^3^H]5-CT-labeled 5-HT7Rs under different experimental conditions

#### 5.1. Test Ligands

Radioligand competition binding assays were performed to evaluate the affinities of 5-CT maleate (Tocris), SB-269970 hydrochloride (Tocris), and lurasidone (MedChemExpress) (Figure 1). Freshly prepared 10 mM stocks of each compound in dimethylsulfoxide (DMSO) were used for each biological test replicate to account for error in weighing. [^3^H]5-CT was used to label 5-HT7Rs (PerkinElmer, lots 2942594 and 3104075) and was prepared each day of testing in one of the two buffers.

**Figure 1.**
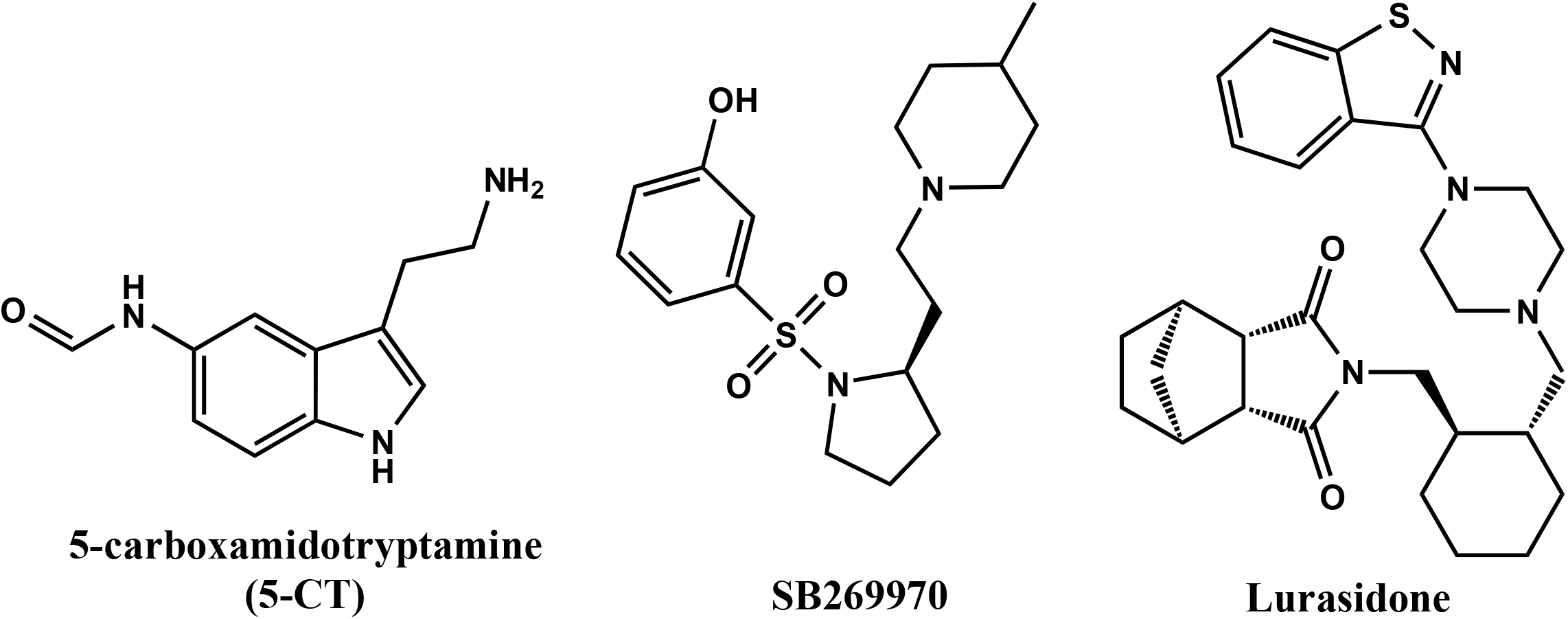
Structures of ligands tested at 5-HT7Rs

#### 5.2. Competition Binding Assays

To assess ligand affinities, competition binding assays were performed under four experimental conditions: SBB at RT, SBB at 37°C, abECF at RT, and abECF at 37°C. Ligands were serially diluted in SBB or abECF from their 10 mM DMSO stock solutions. In 96-well plates, the reaction mixture included either SBB or abECF (150 µL), 0.2 nM final concentration of [^3^H]5-CT (25 µL in buffer), test ligands (25 µL in buffer), and the prepared cell membranes (50 µL in buffer) at RT or 37°C —biological replicates in SBB and abECF were tested by side-by-side on the same day for direct comparisons. Non-specific binding was defined by 10 μM lurasidone. Plates were sealed with transparent tape, wrapped in aluminum foil, and incubated for 90 min at RT or 37°C on a shaker. Following incubation, plate contents were vacuumed to capture bound radioligand on GF/B fiberglass filter mats. The filters were then washed with 250 mL of ice-cold, 50 mM Tris-HCl assay buffer (50 mM Tris-HCl, pH 7.4 at 4°C) to remove unbound radioligand. The wash was repeated twice, and then the filters were dried on a hot plate for approximately 30–45 min. The dry filters were then soaked in plastic bags with Betaplate scintillation fluid, which were sealed and placed in cassettes. Tritium-elicited scintillations were detected and recorded by a Microbeta2 microplate counter, and DPM, corresponding to each well on the plate were recorded for two minutes. Six biological replicates were tested for each experimental condition. For each biological replicate, there were two (5-CT) or three (SB-269970, lurasidone) technical replicates. To explore the contribution of NaCl to the affinities of SB-269970 and lurasidone determined in abECF, a separate experiment was performed in abECF with no NaCl at RT; for these assays, there were two biological replicates, with eight and four technical replicates for lurasidone and SB-269970, respectively; 0.1% bovine serum albumin (fraction V, fatty acid free, EMD Millipore) was included with abECF to also account for the possibility that lurasidone’s affinity was affected by insufficient solubility in SBB or abECF.

## Data analysis

### 1. 5-HT7R saturation binding

DPM from each [^3^H]5-CT concentration were plotted in GraphPad Prism 10. DPM from total binding (TB) were subtracted from DPM from nonspecific binding (NSB) to obtain specific binding, i.e., (SB), SB = TB – NSB, and these values were transformed to fmol/mg of protein. Data were analyzed using a one-site specific binding model, with least-squares regression. Due to the low number of biological replicates, raw values from individual assays testing each experimental condition were combined to facilitate statistical analyses; raw values were also combined to generate graphs. Extra sum-of-squares F-tests were used to evaluate the impact of temperature and buffer on *K*_d_ and B_max_. Alpha was set to 0.05.

### 2. 5-HT7R competition binding assay

DPM for each concentration of test ligand were plotted in GraphPad Prism. Specific binding was determined as described above. Data were analyzed using a one-site fit *K*_i_ model compared to a two-site fit *K*_i_ model. Hill slopes were determined from a log(inhibitor) vs. response, variable slope (four parameters) model. The *K*_d_ was set to 0.2 nM (the mean *K*_d_ value of [^3^H]5-CT determined from saturation binding experiments). The data, expressed as the percentage of [^3^H]5-CT bound at various concentrations, was graphed against the logarithm of the radioligand concentration to determine the half-maximal effective concentration (IC_50_). GraphPad then applied the Cheng-Prusoff equation to calculate the *K*_i_ values (Yung-Chi and Prusoff, 1973). p*K*_i_ values from individual assays, N=6 per experimental condition, were obtained, and one-way ANOVAs were performed to compare the effects of experimental conditions on mean p*K*_i_ values for each ligand. Post-hoc tests were performed using Tukey’s multiple-comparison test. Results from individual assays, testing each experimental condition, were combined to generate a single graph for each condition, which reports mean and SEM.

## Results

### 1. Experimental conditions impacted [^3^H]5-CT–5-HT7R binding site density, but not [3H]5-CT affinity at 5-HT7Rs

We employed saturation binding assays to evaluate the *K*_d_ of [^3^H]5-CT across various experimental conditions, during which we also determined the number of labeled receptor sites B_max_ (Figure 2, Table 2). R^2^ values from non-linear regression fits of results from each biological replicate ranged from 0.93 to 0.99, supporting the one-site specific binding model. *K*_d_ values for each experimental condition—obtained by analyzing results from all technical replicates per condition—were consistent and ranged from 0.21 to 0.31 nM. There were no significant differences in *K*_d_ between conditions. Best-fit B_max_ values ranged from 2,174 to 3,403 fmol/mg protein between the various conditions. There was a trend towards an increase in B_max_ at 37°C, irrespective of buffer (F (1, 188) = 3.021, P = 0.0838). Incubation in abECF significantly increased B_max_ relative to SBB, irrespective of temperature (F (1, 188) = 20.44, P < 0.0001).

**Table 2.** Affinities (*K*_d_) of [^3^H]5-CT at 5-HT7 receptors and 5-HT7 receptor binding site densities (B_max_) across experimental conditions obtained from saturation binding. *K*_d_ values were consistent across experimental conditions and aligned with previously reported *K*_d_ values. Incubation in abECF tended to increase B_max_ relative to SBB.

| [<sup>3</sup>H]5-CT | SBB RT | SBB 37° | abECF RT | abECF 37° |
| --- | --- | --- | --- | --- |
| $K_d$, nM<br>(95% CI) | 0.2284 (0.1120–<br>0.4497) | 0.3134 (0.1809–<br>0.5364) | 0.2049 (0.1482–<br>0.2812) | 0.2258 (0.1501–<br>0.3347) |
| $B_{max}$, fmol/mg<br>(95% CI) | 2174 (1803–<br>2602) | 2682 (2299–<br>3140) | 3170 (2918–<br>3444) | 3403 (3059–<br>3784) |

**Figure 2.**
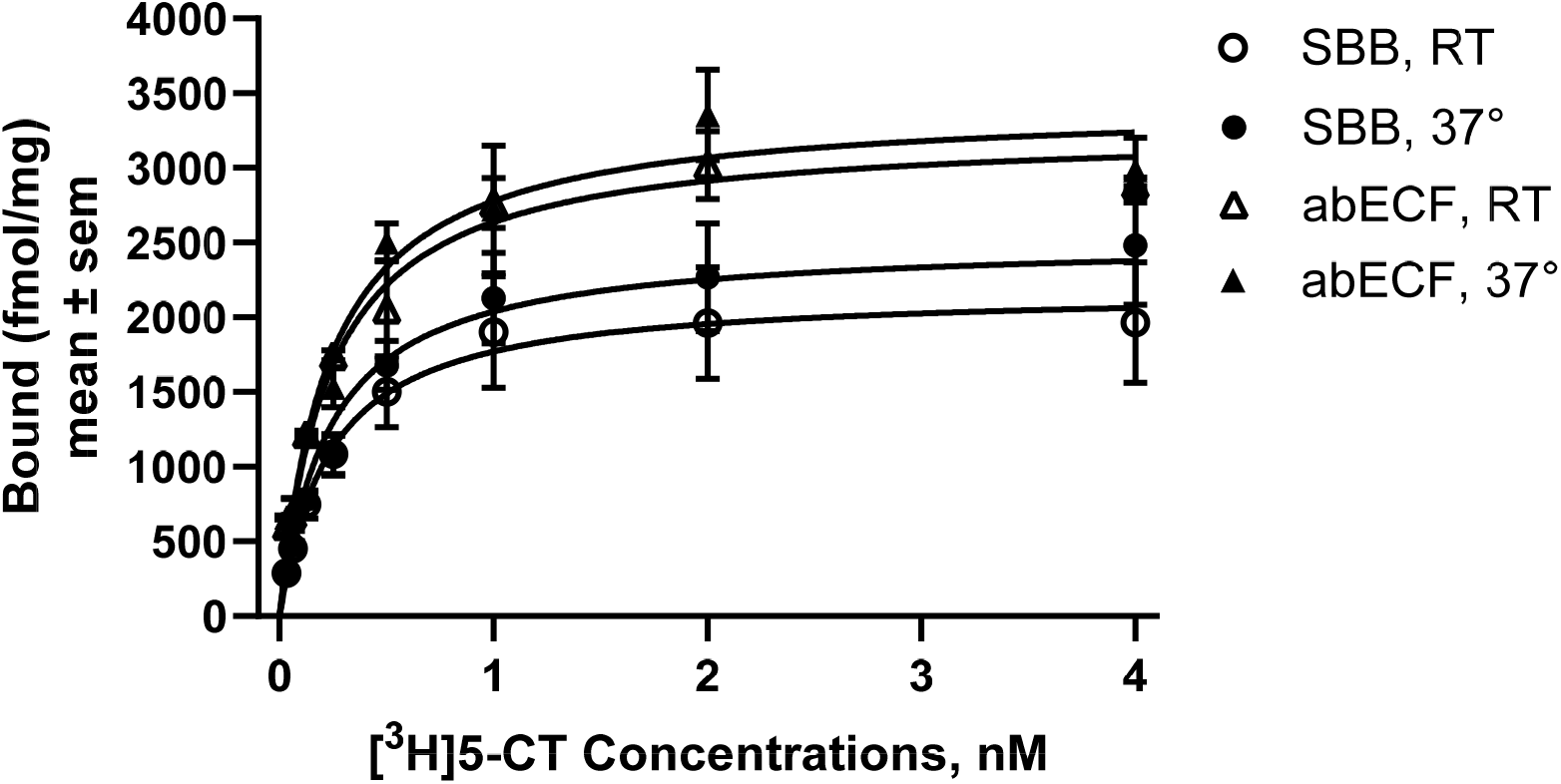
Saturation binding isotherms of [^3^H]5-CT at 5-HT7 receptors across experimental conditions.

### 2. Environmental conditions impacted the binding affinity of antagonists at 5-HT7Rs

Like the *K*_d_ for [^3^H]5-CT, the *K*_i_ for 5-CT determined in the competitive radioligand binding experiment at 5-HT7Rs was not influenced by either the temperature or buffer conditions (Figure 3, Table 3) (F (3, 20) = 1.488, P = 0.2480). However, SB-269970’s affinity was dependent on experimental conditions (F (3, 20) = 23.44, P < 0.0001). Post-hoc analyses revealed an increase in SB-269970’s affinity at 37°C compared to RT when incubated in SBB (P = 0.0031) and an increase in affinity when incubated in abECF compared to SBB at RT (P < 0.0001) and at 37°C (P = 0.0070). Like SB-269970, lurasidone’s affinity was also dependent on experimental conditions (F (3, 20) = 25.59, P < 0.0001). Post-hoc analyses revealed an increase in lurasidone’s affinity at 37°C compared to RT when incubated in SBB (P = 0.02) and an increase in affinity when incubated in abECF compared to SBB at RT (P < 0.0001) and at 37°C (P = 0.0016). R^2^ values from the one-site fit, *K*_i_ model for 5-CT, SB-269970, and lurasidone were: 1) 0.9790, 0.9292, and 0.8844 with SBB at RT; 2) 0.9836, 0.9805, and 0.9700 with SBB at 37°C; 3) 0.9826, 0.9787, and 0.9504 with abECF at RT; and 4) 0.9844, 0.9837, and 0.9641 with abECF at 37°C, respectively. Hill slopes for 5-CT, SB-269970, and lurasidone data were: 1) -1.000, - 0.7827, and -1.007 with SBB at RT; 2) -0.9654, -0.8084, and -0.9692 with SBB at 37°C; 3) -1.011, -0.7991, and -0.6658 with abECF at RT; and 4) -1.001, -0.9361, and -0.7880 with abECF at 37°C, respectively. The Hill slopes that were less than unity for SB-269970 and lurasidone could indicate heterogeneous binding sites for the antagonists or negative cooperativity; indeed, the two-site fit, *K*_i_ model was preferred over the one-site fit, *K*_i_ model for SB-269970 and lurasidone in most conditions (P = 0.005 to < 0.0001) (Figure 4, Table 3). Though, R^2^ values for SB-269970 data were improved by less than 0.01 for the two-site fit for all experimental conditions; R^2^ values for lurasidone improved for the two-site fit by less than 0.002 for SBB conditions, by 0.03 for abECF at RT, and by 0.01 for abECF at 37°C. Removing NaCl from the abECF buffer had a minor affinity-enhancing effect on lurasidone and SB-269970, compared to normal abECF, as determined using a one-site *K*_i_ model.

**Figure 3.**
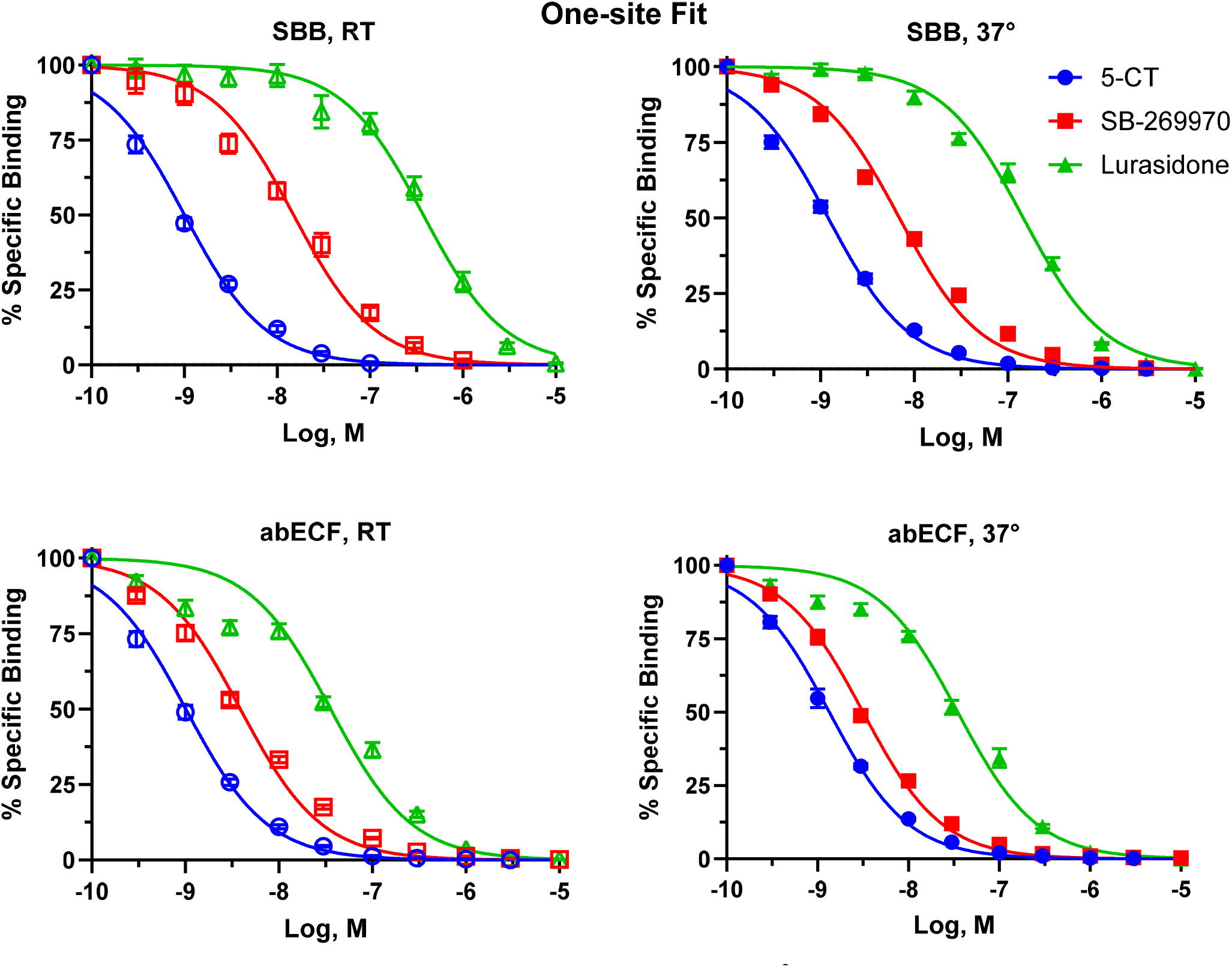
Competition binding results from test ligands at [^3^H]5-CT-labeled 5-HT7 receptors across experimental conditions; data are fit to a one-site fit *K*_i_ model. Note the increased affinity of SB-269970 and lurasidone observed when incubations were performed in abECF (bottom) compared to SBB (top). Physiological temperature also modestly enhanced the affinity of SB-269970 and lurasidone.

**Figure 4.**
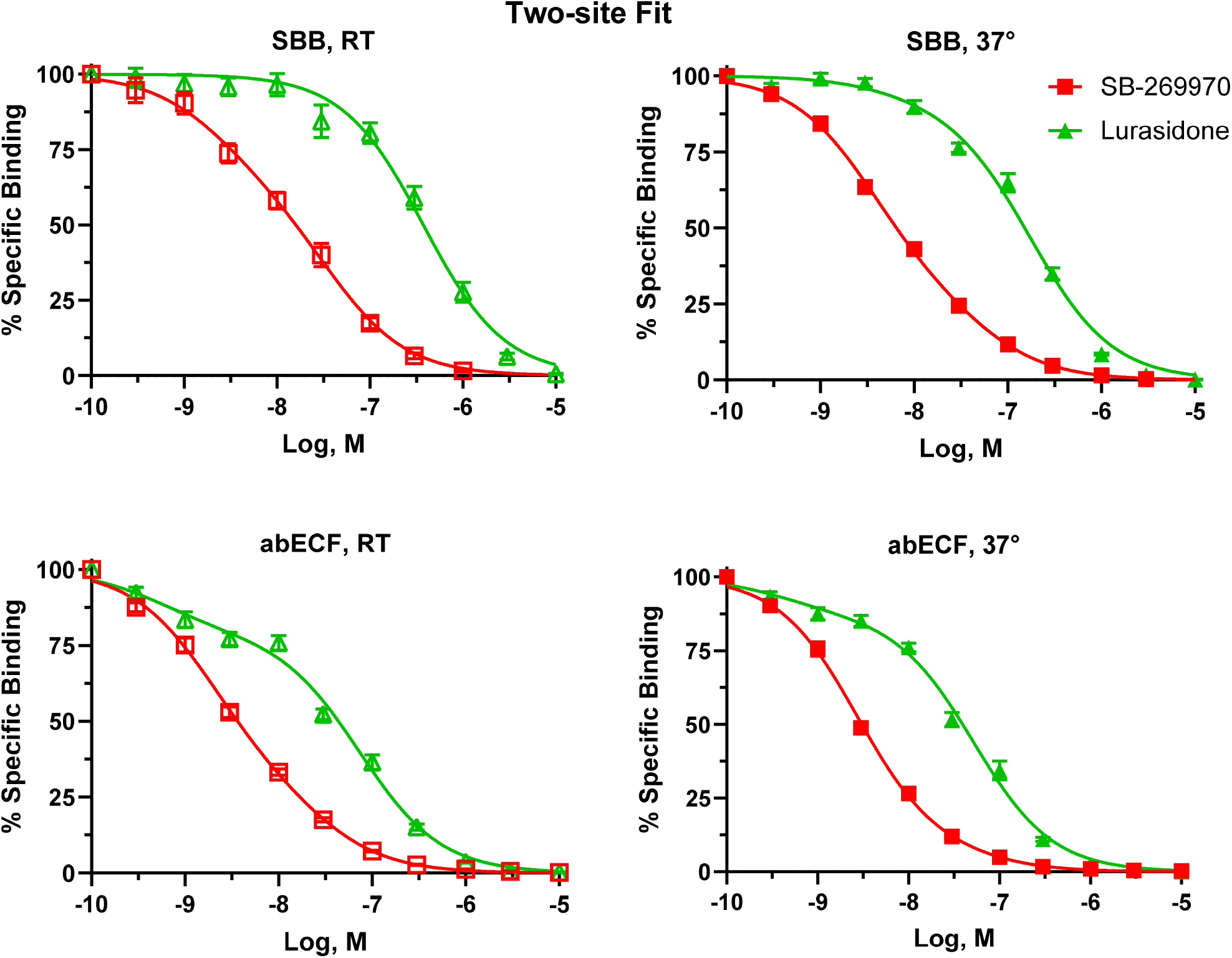
Competition binding results from antagonist ligands at [^3^H]5-CT-labeled 5-HT7 receptors across experimental conditions; data (same as in Figure 3) are fit to a two-site fit *K*_i_ model. Note the increased affinity of SB-269970 and lurasidone observed when incubations were performed in abECF (bottom) compared to SBB (top). Physiological temperature also modestly enhanced the affinity of SB-269970 and lurasidone.

**Table 3.** Affinities of test ligands at 5-HT7 receptors across experimental conditions obtained from [^3^H]5-CT competition binding. Results show *K*_i_ values in nM (p*K*_i_, 95% CIs) from one-site and two-site fit *K*_i_ models. A one-site fit was preferred for 5-CT in all conditions, so only one-site results are shown for 5-CT. Note the temperature and assay buffer-dependent changes in affinities of the antagonists. To assess the contribution of physiological concentrations ofNaCl (128 mM) to antagonist affinities obtained with abECF, separate tests were conducted at RT with abECF containing no NaCl; note that no NaCl had a modest affinity-enhancing effect. N.D. = not determined

| Model | Ligand | SBB RT | SBB 37° | abECF RT | abECF 37° | abECF RT –<br>No NaCl |
| --- | --- | --- | --- | --- | --- | --- |
| One-site | 5-CT | 0.371 (9.430,<br>9.465–9.396) | 0.445 (9.351,<br>9.382–9.321) | 0.374 (9.428,<br>9.459–9.396) | 0.496 (9.304,<br>9.334–9.273) | N.D. |
|  | SB-269970 | 5.55 (8.256,<br>8.320–8.192) | 2.56 (8.592,<br>8.624–8.559) | 1.43 (8.844,<br>8.877–8.812) | 1.17 (8.932,<br>8.959–8.906) | 0.39 (9.412,<br>9.483–9.342) |
|  | Lurasidone | 139 (6.859,<br>6.930–6.787) | 53.9 (7.269,<br>7.308–7.230) | 13.2 (7.880,<br>7.940–7.820) | 12.9 (7.890,<br>7.937–7.843) | 9.27 (8.033,<br>8.117–7.949) |
| Two-site | High<br>SB-269970 | 0.92 (9.036,<br>9.577–8.659) | 1.25 (8.903,<br>9.043–8.800) | 0.67 (9.176,<br>9.329–9.066) | 0.99 (9.004,<br>9.114–8.949) | 0.17 (9.774,<br>9.943–9.641) |
|  | Low<br>SB-269970 | 13.7 (7.863,<br>8.070–7.554) | 16.8 (7.774,<br>8.037–7.458) | 8.81 (8.055,<br>8.308–7.747) | Unstable | 3.27 (8.486,<br>8.797–8.093) |
|  | High<br>Lurasidone | One-site<br>Preferred | 2.22 (8.654,<br>11.01–7.988) | 0.20 (9.701,<br>9.972–9.443) | 0.15 (9.823,<br>10.28–9.393) | 1.38 (8.860,<br>9.982–8.354) |
|  | Low<br>Lurasidone | One-site<br>Preferred | 65.1 (7.186,<br>7.256–7.101) | 27.5 (7.561,<br>7.637–7.481) | Unstable | 29.0 (7.537,<br>7.868–6.939) |

## Discussion

Accurate and precise ligand pharmacology assays are important for guiding structure–activity studies and essential for reliable selectivity profiling (Sum et al., 2019). We thus evaluated the impact of performing radioligand 5-HT7R binding assays at physiological temperature in abECF to better model physiological conditions in the brain where ligands interact with GPCRs expressed in the cell membrane, comparing affinities to those obtained from incubation in abECF at RT and incubation in SBB, at physiological temperature and RT.

### Temperature

Temperature did not affect the affinity of 5-CT at 5-HT7Rs as determined by saturation and competitive radioligand binding, and the 5-CT affinities we observed here aligned with our and others’ previous reports, regardless of assay conditions (Armstrong et al., 2020; Brüss et al., 2005; Eglen et al., 1997; Thomas et al., 1998). Physiological temperature, however, tended to increase [^3^H]5-CT–5-HT7R receptor binding sites (B_max_) relative to RT, suggesting an increase in 5-HT7R active state conformations, as has been observed with other GPCRs (Borea et al., 1996; Thakur et al., 2023). Conversely, physiological temperature increased the affinities of the antagonists lurasidone and SB-269970 at 5-HT7Rs.

The change in free energy (ΔG) upon binding is the difference in free energy between the unbound and the bound state (ligand–receptor complex). It is defined as ΔG = ΔH – TΔS, where ΔH is the enthalpy change, T is the absolute temperature, and ΔS is the entropy change. A negative ΔG (ΔG < 0) indicates that formation of the ligand–receptor complex is thermodynamically favorable and occurs spontaneously under standard conditions. A number of phenomena contribute to the binding affinity, including both enthalpic and entropic contributions. The affinity, or more specifically, the equilibrium dissociation constant (*K*_d_), is defined as the ratio of the dissociation rate constant (*k*_off_) to the association rate constant (*k*_on_): *K*_d_ = *k*_off_ /*k*_on_. Here, *k*_on_ (M^-1^S^-1^) describes how quickly the ligand-receptor complex forms, and *k*_off_ (S^-1^) describes how quickly the ligand-receptor complex dissociates. Temperature could alter affinity by changing either *k*_on_ and *k*_off_ or both, and additional experiments are required to determine such effects for 5-CT, lurasidone, and SB-269970. One limitation of the current experiments is that while we used a relatively standard 90-min incubation time, which appears sufficient for [^3^H]5-CT and 5-CT to reach equilibrium—because the affinity of 5-CT did not depend on temperature—we cannot rule out the possibility that equilibrium was not achieved in all experiments (Armstrong et al., 2020). In other words, the temperature-dependent change in affinity for lurasidone and SB-269970 could be a kinetic phenomenon. Specifically, a higher physiological temperature could shorten the *k*_on_ rate, allowing equilibrium to be reached faster at 37°C and thus appear as a higher affinity at the time points used. Diffusion rates are inversely related to size (as described by the molecular radius in the Stokes-Einstein equation) (Miyamoto and Shimono, 2020). Thus, it is notable that the antagonists lurasidone, 492.68 g/mol, and SB-269970, 352.49 g/mol, are larger than 5-CT, 203.25 g/mol. Also, 5-CT’s logP is -0.31, SB-269970’s is 2.39, and lurasidone’s is 5.25, and logP’s correlated with the increasing impact of temperature (and abECF) on affinity. The larger size and logP values could affect the *k*_on_ rate through various phenomena, including initial receptor engagement and desolvation differences due to the larger solvent accessible surfaces (Sims et al., 2005).

Differences in *k*_off_, resulting from changes in the strength of various ligand-receptor interactions, could also explain the higher affinity. 5-HT7R ligands are known to make a number of ligand-receptor residue interactions within the orthosteric binding site of 5-HT7R. These include identified Van der Waals (e.g., CH−aryl and aryl-aryl) interactions with phenylalanine F6.52, F6.51, or F3.28, and hydrogen bonding with tyrosine Y7.43 or serine S5.43, in addition to ionic interactions between the ammonium cation on virtually all 5-HTR ligands and the highly conserved aspartic acid D3.32 residue (Canal et al., 2011; Impellizzeri et al., 2015; Kołaczkowski et al., 2006; López-Rodr_í_guez et al., 2000; Vermeulen et al., 2004). A recent cryo-electron microscopy (cryo-EM) study determined the structure of 5-CT-bound 5-HT7R–Gα_s_ complex (Huang et al., 2022). In addition to the interactions mentioned previously, isoleucine residue I233 from extracellular loop 2 (ECL2) makes Van der Waals contacts with 5-CT. Active vs. inactive state structures of a similar class A GPCR show conformational changes in the extracellular portions of the receptor, including the extracellular vestibule (Kim et al., 2020). A number of studies have found that agonist-induced activation of the receptor results in a contraction of the extracellular binding pocket; however, it appears this may be ligand-specific (Kim et al., 2020). ECL2 acts as a lid over the orthosteric binding site of many class A GPCRs and can influence ligand binding affinities. For example, the dissociation rate of LSD at the 5-HT2AR and 5-HT2BR has been shown to depend on interactions with an ECL2 leucine, L209. For example, mutation of this residue to alanine leads to 10-fold reduction in residence time (dependent on *k*_off_) at the 5-HT2BR at 37°C. However, interestingly, the mutation reportedly increased the *K*_on_ rate of LSD, so the equilibrium binding affinity was not changed (Kim et al., 2020; Wacker et al., 2017), illustrating the complex multicomponent nature of binding affinity.

It is believed that most orthosteric GPCR ligands initially make contact with receptor amino acid residues on the extracellular surface that comprise what is known as the extracellular vestibule, which includes ECL2. This interaction occurs prior to desolvation and shuttling to the orthosteric binding site (Latorraca et al., 2017). In the current study, one possibility for the compound-specific temperature dependence of affinity is that temperature influences the conformation of solvent-exposed ECL2 residues, including I233, which differentially impacts its interactions across the three compounds. Consistent with this, hydrophobic effects are known to be temperature dependent, and temperature affects protein structure (van Dijk et al., 2015). It could be that these conformational changes at 37°C strengthen interactions (and lengthen the dissociation rate) between the receptor and the larger, more lipophilic antagonists. Another 5-HT2AR agonist, mescaline, which is notably smaller than LSD (molecular weight: 211.26 vs. 323.43 g/mol, respectively) also makes contact with L209, but did not show the same impact on dissociation rate as LSD (Gumpper et al., 2025). Future work could explore the potential role of ECL2 residues, including I233, in governing this effect with 5-HT7 binding affinities via mutagenesis studies.

One final note is that, at dopamine D2Rs, the affinity of the D2R antagonist raclopride was several-fold lower at physiological temperature compared to RT (Hall et al., 1990), suggesting the effects of temperature likely depend on both the ligand and GPCR tested. Performing pharmacology experiments at physiological temperature should be strongly considered, as it mimics the endogenous environment. Most of the current understanding of biomolecular ligand–GPCR interactions is based on high-resolution X-ray crystal structures and those obtained from cryogenic electron microscopy (cryo-EM) (Benjin and Ling, 2020). However, these techniques typically involve cooling samples to cryogenic temperatures, which could alter GPCR conformations from those observed at physiological temperatures. Recent X-ray crystallography research demonstrated different protein conformations, binding sites, and poses when structures were examined at RT compared to cryogenic temperatures (Skaist Mehlman et al., 2023), suggesting that experiments conducted at physiological temperatures could yield insights more closely aligned to biological conditions within living organisms.

### Buffer

Another factor that can impact the affinity of ligands is their interactions with ions in the assay incubation medium. Na^+^ and K^+^ in abECF are essential for neural functions, including establishing cell membrane potentials and generating action potentials. At physiological concentrations, Na^+^ binds to a conserved sodium pocket in most class A GPCRs—interacting with water molecules and anchoring ionically at D2.50—and behaves as a negative allosteric modulator of various class A GPCRs by inhibiting conformational changes in the orthosteric binding pocket and stabilizing inactive conformations, thereby decreasing the affinity of some agonists and increasing the affinity of some antagonists (Draper-Joyce et al., 2018; Hall et al., 1990; Katritch et al., 2014; Limbird et al., 1982; Michino et al., 2015; Neve, 1991; Strasser et al., 2015; van der Westhuizen et al., 2015; Zarzycka et al., 2019). This effect was not evident in our data, as the agonist 5-CT displayed the same affinity when using NaCl-lacking SBB or NaCl-containing abECF, and we observed only a minor increase in antagonist affinity when NaCl was excluded from abECF. In another study, NaCl had no effect on agonist affinity, but increased B_max_ of agonist-labeled histamine H3Rs (Schnell and Seifert, 2010), suggesting Na^+^ might have contributed to the increase in B_max_ we observed with [^3^H]5-CT-labeled 5-HT7Rs incubated in abECF. As is the case with temperature, effects of Na^+^ and other ions on ligand affinities can vary markedly between receptors and between ligands of the same receptor, and is likely due to the specific affinity of ions at their binding sites in the receptors and their unique effects on the conformations of residues at which distinct ligands interact (Zarzycka et al., 2019). While there is little evidence that K^+^ and Cl^-^ influence ligand-receptor binding, high concentrations of divalent ions such as Mg^2+^ and Mn^2+^, but not Ca^2+^, have been found to impact the binding affinity of both agonists and antagonists at GPCRs, e.g., µ-opioid and 5-HT1ARs (Kalipatnapu et al., 2004; Pasternak and Pan, 2013).

abECF includes 1.06 mM glucose, the approximate glucose concentration in brain ECF (de Vries et al., 2003; McNay and Sherwin, 2004). Glucose has been found to act as a positive modulator at the glycine receptor, a cys–loop ion channel receptor (Breitinger et al., 2015), and, in a calcium-dependent manner, is a positive modulator of the calcium-sensing receptor, a class C GPCR (Medina et al., 2016). While there is no current evidence of glucose influencing ligand binding at class A GPCRs, including 5-HT7Rs, this remains a hypothesis worth exploring. Similarly, there appears to be no well-established, specific effects of phosphate buffers on ligand–GPCR interactions; however, phosphate buffers contribute to the overall ionic strength of the solution. One report noted an increase in antagonist-labeled adrenoceptor 2 binding sites when membranes were incubated in 50 mM Na-K phosphate buffer compared to 50 mM Tris-HCl buffer (Perry and U’Prichard, 1981). The impact of the phosphate buffer in abECF is unknown, but given that it is a physiological buffer found in CNS ECF, it should be evaluated further. Ionic strength reflects the concentration and charge of ions in solution. Ionic strength can affect protein conformation by altering the balance of electrostatic interactions within the protein itself or between the protein and the surrounding solvent (Berg et al., 2023). Changes in receptor conformation can, in turn, alter the accessibility of the ligand binding site, thereby affecting the binding affinity of a given ligand. Thus, the difference in the ionic strength of the buffers may have differently impacted antagonist binding at 5-HT7Rs.

There is considerable heterogeneity in the compositions of incubation media used in ligand–class A GPCR binding assays. At 5-HT7Rs, besides those mentioned in the Introduction, other media include 50 mM Tris HCl, 4 mM MgCl_2_, 10 mM pargyline, and 0.1% ascorbate (Kucwaj-Brysz et al., 2024). The following provides other examples from class A GPCR ligand binding studies. At dopamine receptors, media used include Tris-HC1 buffer containing 0.1% ascorbic acid, 120 mM NaCl, 5 mM KCl, 2 mM CaC1_2_, and 1 mM MgC1_2_ (Hall et al., 1990) and 170 nM Tris HCI buffer containing 120 mM NaCI, 5 mM KCl, 1.5 mM CaCl_2_, 4 mM MgC1_2_, and 1 mM EDTA (Ricci and Amenta, 1994). At histamine receptors, media include 50 mM Tris-HCI containing 2 mM MgCl_2_ (Moguilevsky et al., 1994) and 50 mM Na_2_HPO_4_ with 50 mM KH_2_PO_4_ (Gbahou et al., 2006). At adrenoceptors, media include 50 mM Tris HCl with 1 mM EDTA (Oshita et al., 1991) and 50 mM Na-K phosphate buffer or 50 mM Tris-HCl (Perry and U’Prichard, 1981). At muscarinic receptors, media include phosphate buffered saline with 5 mM EDTA (Peralta et al., 1987), 0.03 mM HEPES buffer containing 0.5 mM MgCl_2_ and 0.5 mM EGTA (Christopoulos and Wilson, 2001), 50 mM HEPES containing 110 mM NaCl, 5.4 mM KCl, 1.8 mM CaCl_2_,1mM MgSO_4_, 25 mM glucose and 58 mM sucrose (Christopoulos et al., 1998), 20 mM HEPES buffer containing 1 mM MgCl_2_ (Cembala et al., 1998), and Krebs-Henseleit buffer containing 118 mM NaCl, 4.7 mM KC1, 1.2 mM MgSO_4_, 1.3 mM CaC1_2_, 25 mM NaHCO_3_, and 1.2 mM glucose (Wang and el-Fakahany, 1993).

These observations illustrate the general lack of consensus on incubation media for evaluating ligand affinities at class A GPCRs; however, Tris buffer tends to be more widely used than sodium phosphate buffers. Tris buffer is often preferred due to its buffering range and versatility across biological applications, e.g., it is also suitable for gel electrophoresis to separate nucleic acids and proteins. Tris has a p*K*_a_ of approximately 8.1 at 25°C, making it an excellent buffer in the physiological pH range of 7.0-9.0. Also, Tris only weakly complexes with Ca^2+^ and Mg^2+^ (Abe et al., 1985; Ferreira et al., 2015; Fischer et al., 1979). This is crucial in enzymatic reactions where these ions are cofactors and their availability needs to be maintained. Despite the advantages of Tris, sodium phosphate buffers are still commonly used. Germane to our rationale, phosphate (unlike Tris) is a naturally occurring physiological buffer in cells and extracellular fluids, making it relevant for mimicking in vivo conditions, and also has excellent buffering capacity at pH 7.4 due to the p*K*_a_ of H_2_PO_4_/HPO_4_^-^.

One complexity in mimicking physiological conditions *in vitro* is that radioligand binding experiments are almost exclusively performed in isolated membrane preparations. In native cells, the extracellular and intracellular regions of membrane-embedded GPCRs will experience very different physiological environments, including ion composition, membrane potential, pH, and associated proteins and other biomolecules. Furthermore, these conditions can be highly dynamic, changing moment to moment. While performing whole-cell radioligand binding may seem like an obvious way to at least partially control for this, it presents its own set of challenges. For one, there is an extensive population of GPCRs inside most cells. In addition, binding often utilizes recombinant receptor overexpression in transfected cells. GPCR trafficking involves moving between intracellular sites (e.g., the endoplasmic reticulum) and the membrane, which is also highly dynamic. The membrane permeability of the radioligand versus the test ligand can differ, resulting in misleading findings. Ultimately, the utility of in vitro experiments depends on the experimental question or technological needs of a given experiment or project. We have shown that the selection of buffer and choice of temperature differentially impacted the binding affinities of three ligands for 5-HT7R.

Presumably, using media that are more proximal to the physiological environment where ligands interact with receptors would lead to more physiologically accurate affinities. It could be broadly adopted to lead to higher reproducibility of affinity values across laboratories (Sum et al., 2019). However, it remains to be seen if this goal is possible. Notably, despite the effects of temperature and buffer on ligand affinities at 5-HT7Rs, the rank order of ligand potencies remained constant across experimental conditions, 5-CT > SB-269970 > lurasidone. Also notable is that the 5-HT7R affinity of lurasidone we observed was substantially lower than previous reports (Horisawa et al., 2013; Ishibashi et al., 2010), regardless of the assay conditions. The reason that the antagonist/inverse agonist binding results fit better to a two-site binding model regardless of assay conditions is unresolved. One possibility relates to the functional pharmacology of the ligands. Assuming that inverse agonists bind to and stabilize the inactive 5-HT7R conformation and also block access of the agonist to the active conformations, it may be that they deplete the population of active site receptors available for agonist binding (e.g., like non-hydrolysable GTP). Thus, inverse agonists might directly compete with agonist radioligands for the active sites with one potency, and also remove receptors available for agonist radioligand binding via shifting the equilibrium away from the active site with another potency; these two potencies would be revealed as two sites in binding curves.

## Conclusions

Incubating ligands and 5-HT7R-expressing cell membranes in abECF at physiological temperature was performed to more closely mimic the in vivo brain environment where ligands bind to cell membranes expressing GPCRs. Our observations demonstrated an increase in the binding affinity of antagonists at 5-HT7Rs when incubated in abECF compared to SBB. Increasing the incubation temperature from RT to 37°C also increased agonist-labeled receptor binding sites. The study emphasizes the importance of comprehensive in vitro modeling that considers the complex biological milieu in which drugs operate.

## Acknowledgments

C.E.C conceived experiments. A.T.H. and H.M.A. performed experiments. A.T.H. and C.E.C. analyzed data. A.T.H, C.E.C., and J.V.W. wrote the manuscript. This work was supported by Mercer University.

